# Interplay Between Retroviral Element De-repression and Wnt/β-Catenin Pathway in Cellular Models of Parkinson’s Disease

**DOI:** 10.64898/2026.02.06.704334

**Authors:** Ewa K. Krasnowska, Sabrina Romanò, Giulia Di Marcantonio, Simona Rossi, Mauro Cozzolino, Annalucia Serafino

**Affiliations:** Institute of Translational Pharmacology, National Research Council of Italy (CNR), Rome, Italy

**Keywords:** Parkinson’s disease, Human endogenous retrovirus K (HERV-K), Wnt/β-catenin pathway, Neurotoxic stress, Neurodegeneration

## Abstract

Parkinson’s disease (PD) is characterized by progressive dopaminergic neurodegeneration driven by complex interactions among oxidative stress, impaired survival signaling, and protein homeostasis disruption. Emerging evidence suggests that endogenous retroelements, including human endogenous retrovirus K (HERV-K), may contribute to neurodegenerative processes; however, their role in PD remains poorly defined. Here, we investigated whether dopaminergic neurotoxic stress induces HERV-K activation and whether modulation of pro-survival signaling pathways influences this response in PD-relevant cellular models. Using undifferentiated SHSY5Y cells and neuron-like retinoic acid–differentiated SHSY5Y cells, we show that exposure to the dopaminergic neurotoxin 6-hydroxydopamine (6-OHDA) induces a rapid and robust transcriptional de-repression of HERV-K Env gene. HERV-K activation occurs early after toxin exposure, scales with the intensity of the insult, and is associated with alterations in oxidative stress defenses, survival signaling pathways, and protein homeostasis. Notably, 6-OHDA treatment promotes the accumulation and cytoplasmic mislocalization of phosphorylated TAR DNA-binding protein 43 (pTDP-43), a pathological feature linked to neurodegenerative proteinopathies. Pharmacological modulation of the Wnt/β-catenin pathway by the natriuretic peptide atrial natriuretic peptide (ANP) significantly attenuates neurotoxin-induced HERV-K activation, restores oxidative stress–related and survival signaling markers, and limits pTDP-43 accumulation and mislocalization. These findings indicate that reinforcement of Wnt/β-catenin dependent protective pathways constrains stress-driven HERV-K de-repression and associated molecular alterations. Overall, this study identifies HERV-K activation as an early stress-responsive feature in PD-like cellular models and supports the existence of a functional interplay between retroelement regulation, survival signaling, and protein homeostasis. Modulation of Wnt/β-catenin signaling may represent a strategy to limit retroelement-associated pathological responses in PD.

## INTRODUCTION

Parkinson’s disease (PD) is a common neurodegenerative disease (ND) characterized by progressive loss of dopaminergic (DA) neurons in the *substantia nigra pars compacta*, together with Lewy body pathology, neuroinflammation, oxidative stress, and mitochondrial dysfunction[1-6]. Despite decades of research, the precise mechanisms that initiate or drive neuronal degeneration in PD remain incompletely understood. About 5% of PD cases are familial forms, which have been associated with autosomal dominant or recessive mutations in SNCA, LRRK2, PINK1, VPS35, Parkin, and DJ-1 genes [7]. The vast majority of cases are sporadic, for which a multifactorial etiology involving environmental insults, genetic predispositions, aging, and dysfunction of key intracellular signaling pathways has been proposed [8-10]. Currently, there is no cure for PD, and no therapy has yet been conclusively demonstrated to modify the underlying disease course. Therapeutic strategies are therefore symptomatic and aim at improving motor and non-motor features, alleviating disability, and enhancing quality of life. The mainstay of pharmacological treatment remains levodopa, often administered in combination with a dopa decarboxylase inhibitor to enhance central bioavailability and minimize peripheral conversion [11]. Levodopa effectively replenishes striatal dopamine and ameliorates cardinal motor symptoms such as bradykinesia, rigidity, and tremor. However, long-term use is associated with complications, including motor fluctuations and levodopa-induced dyskinesias [11, 12]. Thus, the pursuit of novel therapeutic approaches is imperative, and a deeper understanding of the molecular mechanisms underlying the onset of PD is essential for identifying innovative targets that may enable the development of more effective disease-modifying treatments.

One emerging area of interest in ND, and in particular in PD pathogenesis, is the role of Human Endogenous Retroviruses (HERVs), which are remnants of ancient germ-line retroviral infections that now constitute roughly 8% of the human genome [13]. Over prolonged evolutionary timescales, the majority of HERV sequences become transcriptionally silenced due to the accumulation of mutations that result in the loss of their coding capacity [14], or are repressed by cellular epigenetic mechanisms that preserve genomic stability and prevent the activation of potentially harmful genes [15]. However, some HERV families, in particular HERV-K (HML-2), are notable because they retain relatively intact open reading frames and can be transcriptionally reactivated under certain pathological stressors, generating functional transcripts and proteins [16]. Defective repression of endogenous retroviruses (ERVs) has been linked to the development of cancer [17], as well as a range of autoimmune [18] and inflammatory [19] diseases. Recent studies have also revealed a correlation between the de-repression of HERV genes and several neurodegenerative diseases (NDs), including multiple sclerosis, amyotrophic lateral sclerosis, and Alzheimer’s disease [20, 21]. Indeed, several HERV elements exhibit increased expression in the brains of individuals affected by tauopathies and amyotrophic lateral sclerosis (ALS) [22-24]. The initial inflammation of small brain regions, triggered by environmental factors such as diet, chemical pollutants, viral infections, or other exogenous cerebral insults, may activate previously silent HERVs and contribute to neurodegenerative mechanisms. Furthermore, it has been proposed that reactivation of endogenous retroviruses can promote pathological aggregation and spreading of TDP-43 and Tau, contributing to the progression of NDs such as ALS and tauopathies [25, 26], and the targeting of HERV elements has been proposed as a novel therapeutic strategy for ND treatment [27]. Yet, clear evidence of such an association is still lacking for PD.

The Wnt/β-catenin pathway is an evolutionarily conserved molecular signaling cascade that plays an essential role not only in normal embryonic development but also in the maintenance of adult tissues. In the central nervous system (CNS), this pathway regulates multiple aspects of neuronal function, including differentiation, synapse formation, neurogenesis, and neuroprotection [28-30]. Increasing evidence indicates that dysregulation of this pathway is closely associated with the onset of PD and other NDs, such as Alzheimer’s disease (AD) and ALS, and several signaling components have been investigated as potential therapeutic targets [28, 31-34]. Pharmacological (using both synthetic and natural compounds) and cell-based therapies, aimed at activating or modulating this signaling pathway in neurons and glial cells, have demonstrated neuroprotective effects, promoting endogenous neurogenesis and enhancing neuro-restoration in experimental models of PD [10, 31, 35, 36].

Although a correlation between the de-repression of HERV sequences and the dysregulation of the Wnt/β-catenin pathway in PD and other NDs remains to be elucidated, *in vitro* studies suggest that this signaling cascade may act as an upstream regulator of HERV sequences under neoplastic conditions [37]. In this regard, we have previously demonstrated that the natriuretic peptides Atrial Natriuretic Peptide (ANP), Brain Natriuretic Peptide (BNP), and C-type Natriuretic Peptide (CNP), cardiac and vascular-derived hormones also widely expressed in mammalian CNS, can affect the Wnt/β-catenin and, by upregulating this signaling, act as neuroprotective agents in *in vitro* models of PD [35, 36].

In this study, we explored whether the reactivation of HERV-K sequences might contribute to the onset of PD, using *in vitro* models of dopaminergic neurodegeneration. We also examined whether this reactivation is influenced by the modulation of the Wnt/β-catenin signaling pathway under ANP treatment. To this end, two well-established and previously characterized SH-SY5Y cell models— undifferentiated (SH-SY5Ywt) and differentiated (SH-SY5Ydiff) cells—were employed. Both models were exposed to 6-hydroxydopamine (6-OHDA), a neurotoxin that reproduces key features of dopaminergic neuron degeneration observed in PD.

## MATERIALS AND METHODS

### Cell cultures and treatments

SHSY5Y cell line was provided by the American Type Culture Collection (ATCC, Manassas, VA, USA) and was validated by the ATCC cell bank. Exponentially growing SHSY5Y (SHSY5Ywt; Model A) cells were grown as previously described [35], and detailed in **Supplementary Materials and Methods (M&M)**. The differentiated phenotype (SHSY5Ydiff, Model B) was obtained based on protocols previously described [35], and schematized in **Fig. 1A**. Both proliferating (SHSY5Ywt) and retinoic acid(RA)-differentiated (SHSY5Ydiff) SH-SY5Y cells have been previously characterized by our group [35] in terms of morphology and phenotype, and shown to display key features of dopaminergic neurons, thereby representing suitable models for investigating neurotoxicity and neuroprotection in experimental PD. SHSY5Ywt cells express neuronal and dopaminergic markers under basal conditions, whereas SHSY5Ydiff display a more mature neuron-like phenotype, characterized by a block of proliferation, enhanced neurite outgrowth (**Fig. 1A**), and increased expression of dopaminergic markers, making this model more representative of mature dopaminergic neurons [35].

**Figure 1.**
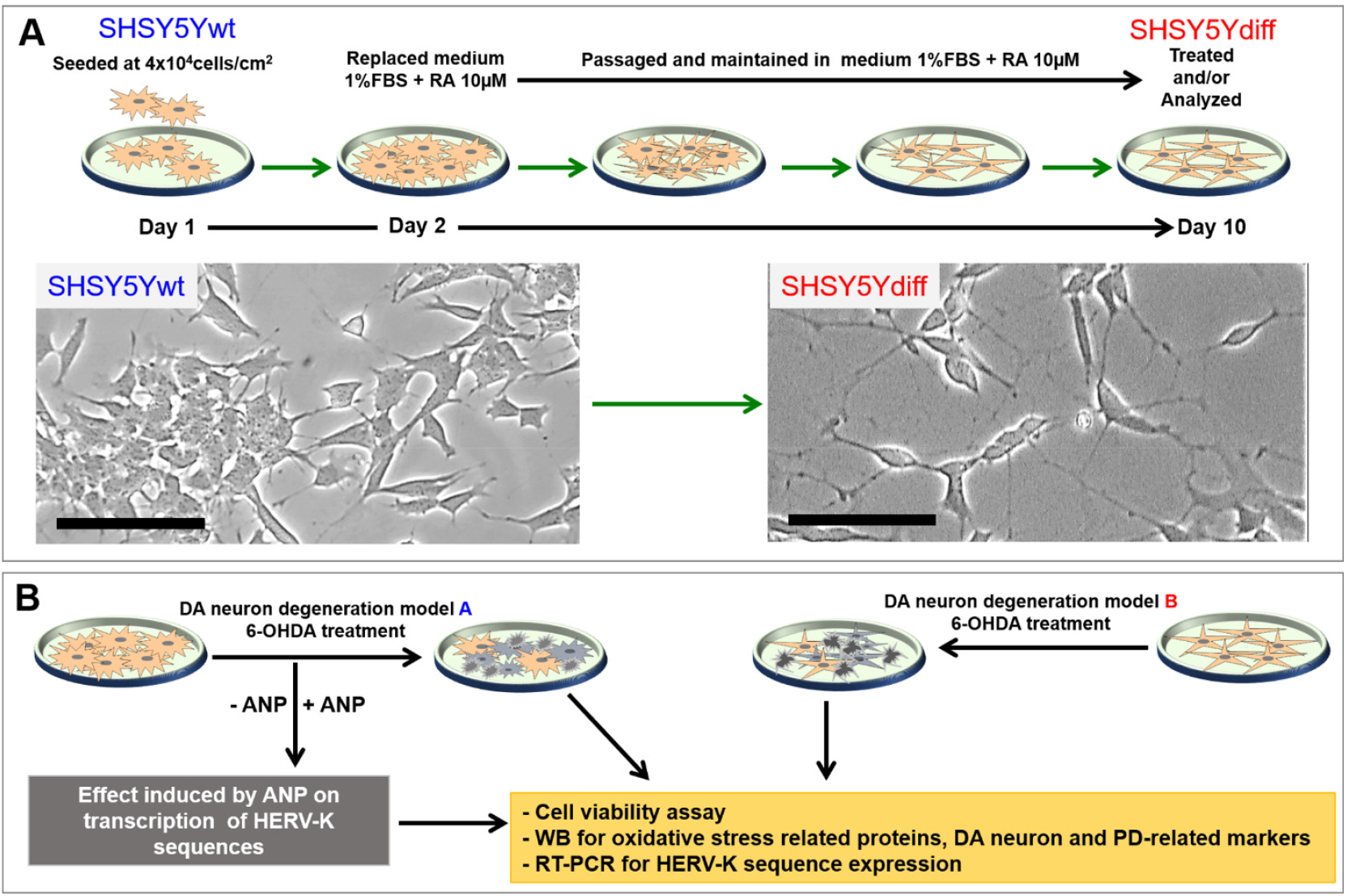
Experimental design and cellular models used in the study. **(A)** Schematic representation of the differentiation protocol of SHSY5Y cells. Undifferentiated SHSY5Y cells (SHSY5Ywt) were seeded at 4 × 10^4^ cells/cm^2^ and subsequently differentiated by culture in low-serum medium (1% FBS) supplemented with retinoic acid (RA, 10 µM) for 9 days, generating neuron-like differentiated cells (SHSY5Ydiff). Representative phase-contrast images illustrate morphological differences between SHSY5Ywt and SHSY5Ydiff cells. Scale bars: 100 µm. **(B)** Schematic overview of the experimental models of dopaminergic neurodegeneration. SHSY5Ywt (Model A) and SHSY5Ydiff (Model B) cells were exposed to 6-hydroxydopamine (6-OHDA) in the presence or absence of atrial natriuretic peptide (ANP). Cells were analyzed for cell viability, oxidative stress– related proteins, dopaminergic and Parkinson’s disease–related markers, phosphorylated TDP-43 localization, and HERV-K sequence expression by RT-qPCR.

For mimicking the neurodegenerative features of PD, SHSY5Ywt and SHSY5Ydiff cells were exposed for 18 h to 25 or 50 µM (for SHSY5Ywt) and 75 µM (for SHSY5Ydiff) of 6-OHDA (prepared in 0.1% ascorbic acid in DMSO), concentrations selected in our previous dose-response studies [35, 36] (**Fig. 1B**). For assessing the ability of the natriuretic peptide in affecting the reactivation of HERV-K sequences induced by the neurotoxic insult, SHSY5Ywt cells were pre-incubated with 100 nM ANP (Phoenix Pharmaceuticals Inc., CA, USA) 30 min before the addition of 6-OHDA to cell cultures, and analyzed up to additional 6 h (**Fig. 1B**, *left panel*).

### Evaluation of Cell Morphology and Viability

Cell morphology was analyzed by phase-contrast microscopy, using the Motic AE31 Trinocular inverted microscope (Motic Asia, Hong Kong). Cell viability was evaluated by the Trypan blue dye exclusion method, followed by analysis using an automated CytoSmart cell counter (Corning, Glendale, AZ, USA).

### Western Blot (WB) Analysis

Cells were lysed in RIPA buffer as detailed in **Supplementary M&M**, and protein concentrations were measured using the Bradford assay (Bio-Rad, Segrate, Italy). Equal amounts of protein (15–20 µg) were resolved by SDS–PAGE on 10% or 15% polyacrylamide gels, transferred onto nitrocellulose membranes (Hybond, Amersham GE Healthcare), and processed for protein immunodetection as detailed in **Supplementary M&M** using the primary antibodies reported in **Supplementary Table S1**. Primary antibody binding was detected using peroxidase-conjugated secondary antibodies (Bio-Rad). Membranes were visualized and images acquired using the ChemiDoc XRS+ imaging system (Bio-Rad). Band intensities were quantified using ImageJ (http://rsbweb.nih.gov/ij/). Densitometric values, normalized to β-actin, were expressed as uncalibrated optical density (OD) values or as fold change relative to untreated controls. Data represent at least three independent experiments and are reported as mean ± SD.

### Quantitative Reverse Transcription Polymerase Chain Reaction (RT-qPCR)

For assessing the transcription levels of HERV-k Env and Gag genes by RT-qPCR, total RNA was extracted and processed for RT-qPCR as detailed **in Supplementary M&M**. The sequences of primers used are reported in **Supplementary Table S2**. Cq values were determined from the system software using ‘single threshold’ mode. The relative expression level of each gene was calculated from these Cqs using experimentally determined amplification efficiencies and then normalized to the reference gene GAPDH. Results were reported as fold of induction *vs* untreated control. In all experiments, each sample was analyzed in triplicate, and no-template controls and no-reverse transcription controls were included.

### Immunocytochemical Analysis and Confocal Microscopy (CLSM)

For confocal microscopy analysis of the intracellular distribution of pTDP-43 aggregates, cells were grown on the ibiTreat µ-Slide 4 wells (Ibidi GmbH, Germany). Immunocytochemical analysis was performed on cells fixed with 4% paraformaldehyde (Sigma–Aldrich). After permeabilization with 0.2% Triton X-100 (Sigma-Aldrich), immunofluorescence staining was done using the primary antibody against pTDP-43 reported in **Supplementary Table S1**. Primary antibody was revealed with Alexa Fluor 488-conjugated anti-mouse or anti-rabbit IgG (Molecular Probes, Eugene, OR, USA). Cell nuclei were counterstained with 0.2% Hoechst (Sigma-Aldrich). Samples were analyzed using the LEICA TCS SP5 confocal microscope (Leica Instruments, Heidelberg, Germany). Quantitative assessment of the percentage of cells exhibiting cytoplasmic pTDP-43 aggregates was performed using ImageJ software. A minimum of 300 cells was analyzed for each sample. Data are presented as the mean ± SD from three independent experiments.

### Statistical analysis

Statistical analyses were conducted using two-tailed Student’s t-tests or one-way analysis of variance (ANOVA), followed by Tukey’s post-hoc test to evaluate differences between groups. A p-value < 0.05 was considered statistically significant. All data derive from at least three independent experiments and are reported as means ± SD.

### Use of AI-assisted editing

Portions of the manuscript text were edited for clarity and readability with the assistance of an artificial intelligence–based language model (ChatGPT, OpenAI). This support was limited to language and stylistic refinement. All scientific content, data interpretation, and conclusions were critically reviewed, validated, and approved by the authors, who take full responsibility for the accuracy, originality, and integrity of the work.

## RESULTS

### Preliminary assessment of neurotoxin-induced de-repression of HERV-k sequences in PD Model A (SHSY5Ywt)

To preliminarily assess whether neurotoxic insult is associated with transcriptional activation of HERV-K sequences, SHSY5Ywt cells were exposed to 6-OHDA for 18 h and compared with cells treated for 18 h with retinoic acid (RA), which was used as a positive control for HERV de-repression, as it has been reported to transiently de-repress specific endogenous retroviral loci in embryonal carcinoma cells [38]. RT-qPCR analysis revealed that exposure to 25 µM 6-OHDA induced a significant increase in the transcription of HERV-K *env* and *gag* sequences relative to untreated controls, with levels comparable to those observed following RA treatment (**Fig. 2**). These data provide initial evidence that dopaminergic neurotoxic stress is sufficient to induce de-repression of HERV-K Env and Gag sequences in SHSY5Ywt cells. We then focused on HERV-K Env transcript because its protein product has been shown to exert neurotoxic effects in human neurons, causing neurite retraction and cell death *in vitro* and neurodegenerative phenotypes *in vivo* [39], and sequences from the Env region have also been implicated in activation of innate immune receptors linked to neuroinflammation [40].

**Figure 2.**
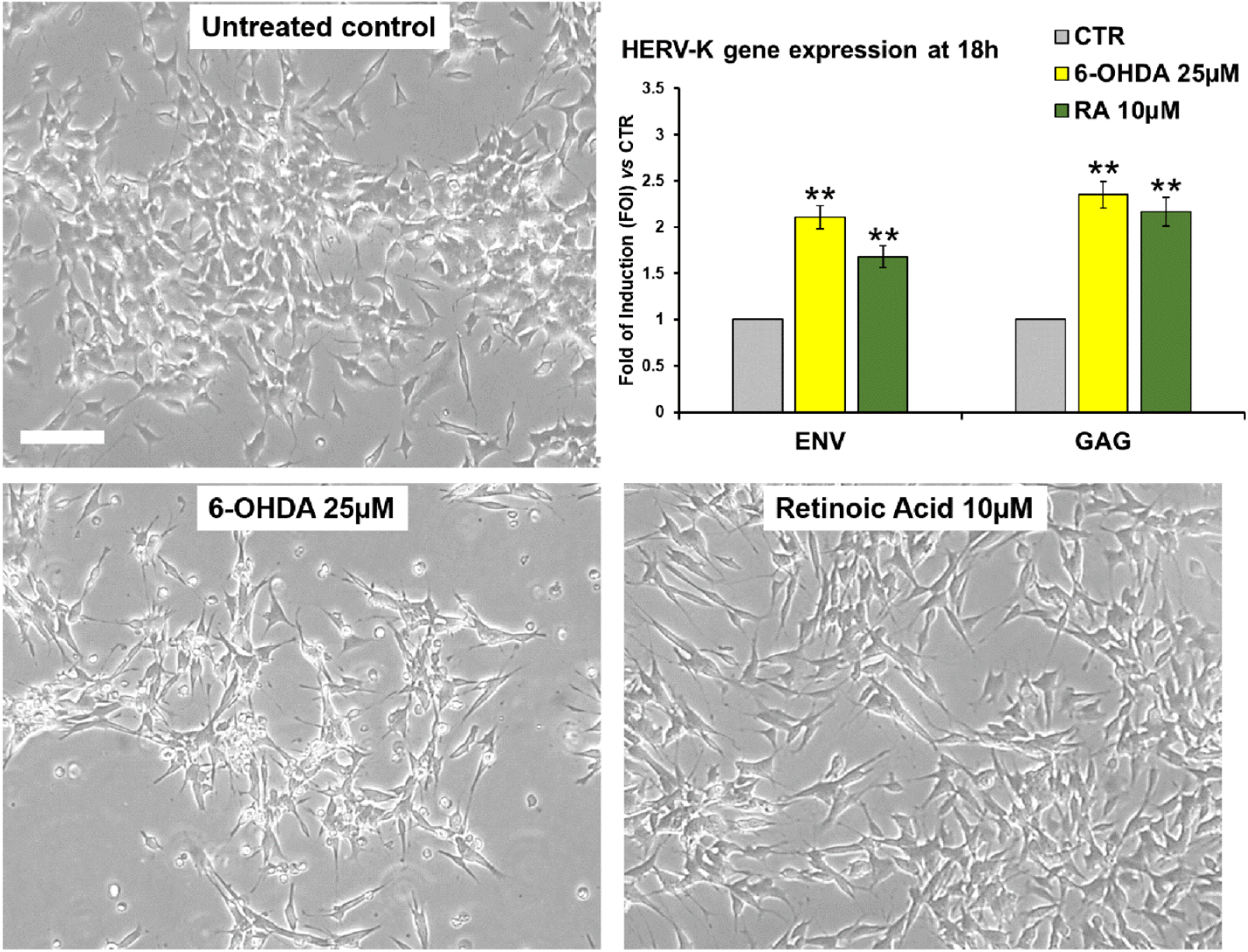
Neurotoxic insult induces transcriptional de-repression of HERV-K sequences in SHSY5Ywt cells. Representative phase-contrast images of SHSY5Ywt cells under untreated conditions or following 18 h exposure to 25 µM 6-OHDA or 10 µM retinoic acid (RA), used as a positive control for HERV de-repression. Scale bars: 100 µm. *Bar graph*: RT-qPCR analysis of HERV-K env and gag gene expression following 18 h treatment. Data are expressed as fold of induction (FOI) relative to untreated controls. Bars represent mean ± SD from at least three independent experiments performed in triplicate. **p < 0.01 *vs* control.

Supporting the hypothesis that the neurotoxic effects of 6-OHDA may involve the reactivation of silent or minimally expressed genomic sequences, washout (WO) experiments demonstrated that the cytotoxic effect is irreversible under the conditions tested (**Fig. 3**). SH-SY5Ywt cells were exposed to 25 or 50 μM 6-OHDA for 18 h and subsequently maintained in neurotoxin-free medium for an additional 24 h. Measurements of monolayer confluence (**Fig. 3B**) showed that, at the end of the 18 h treatment, both 6-OHDA concentrations reduced confluence to approximately 70% of the untreated control. After 24 h of WO, confluence further declined to approximately 30% and 25% for the 25 μM and 50 μM treatments, respectively. Cell viability assessment performed 24 h after WO (**Fig. 3C**), together with optical microscopy analyses (**Fig. 3A**), confirmed that the cytotoxic effect persists following removal of the neurotoxin and displays a clear dose-dependent pattern.

**Figure 3.**
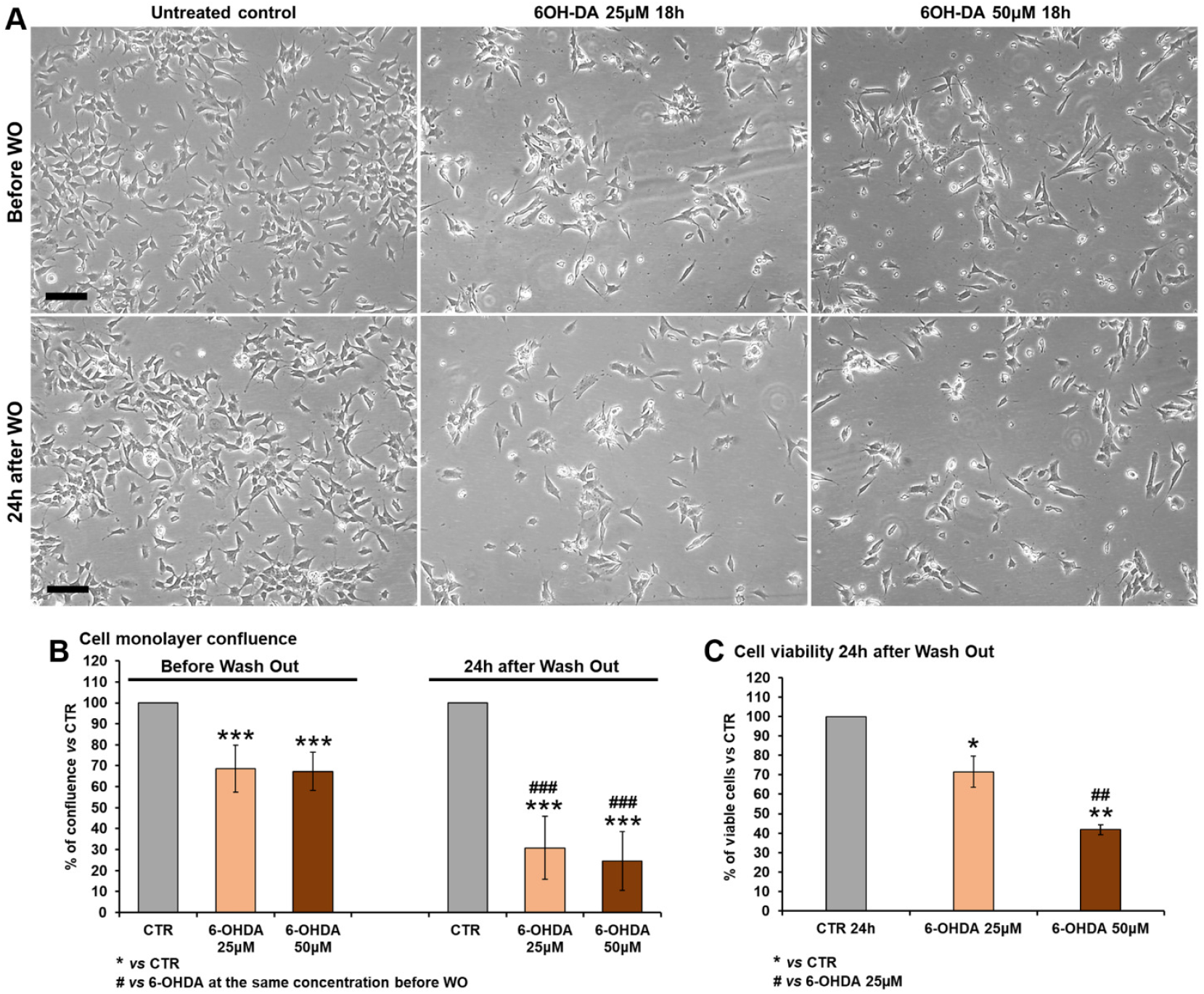
Irreversibility of 6-OHDA–induced neurotoxicity in SHSY5Ywt cells. **(A)** Representative phase-contrast microscopy images of SHSY5Ywt cells before wash-out (WO) and 24 h after removal of 6-OHDA. Cells were treated for 18 h with 25 µM or 50 µM 6-OHDA and subsequently maintained in toxin-free medium for an additional 24 h. Untreated cells are shown as controls. Scale bars: 100 µm. **(B)** Quantitative analysis of cell monolayer confluence before wash-out and 24 h after wash-out. Data are expressed as a percentage of confluence relative to untreated controls. **(C)** Cell viability assessed 24 h after wash-out by Trypan Blue exclusion assay and expressed as a percentage of viable cells relative to control. Data represent mean ± SD from at least three independent experiments. *p < 0.05, **p < 0.01, ***p < 0.001 vs control; #p < 0.05, ##p < 0.01, ###p < 0.001 vs the same 6-OHDA concentration before wash-out.

### 6-OHDA induces dose-dependent neurotoxicity and early HERV-K de-repression in SHSY5Ywt cells

To further characterize the relationship between neurotoxic stress, cellular damage, and HERV-K activation, SHSY5Ywt cells were exposed to increasing concentrations of 6-OHDA. Phase-contrast microscopy revealed marked, dose-dependent morphological alterations after 18 h of treatment, including loss of neuritic extensions, cell rounding, membrane blebbing, and accumulation of cellular debris, which were more pronounced at 50 µM compared to 25 µM 6-OHDA (**Fig. 4A** and **Supplementary Fig. S1**).

**Figure 4.**
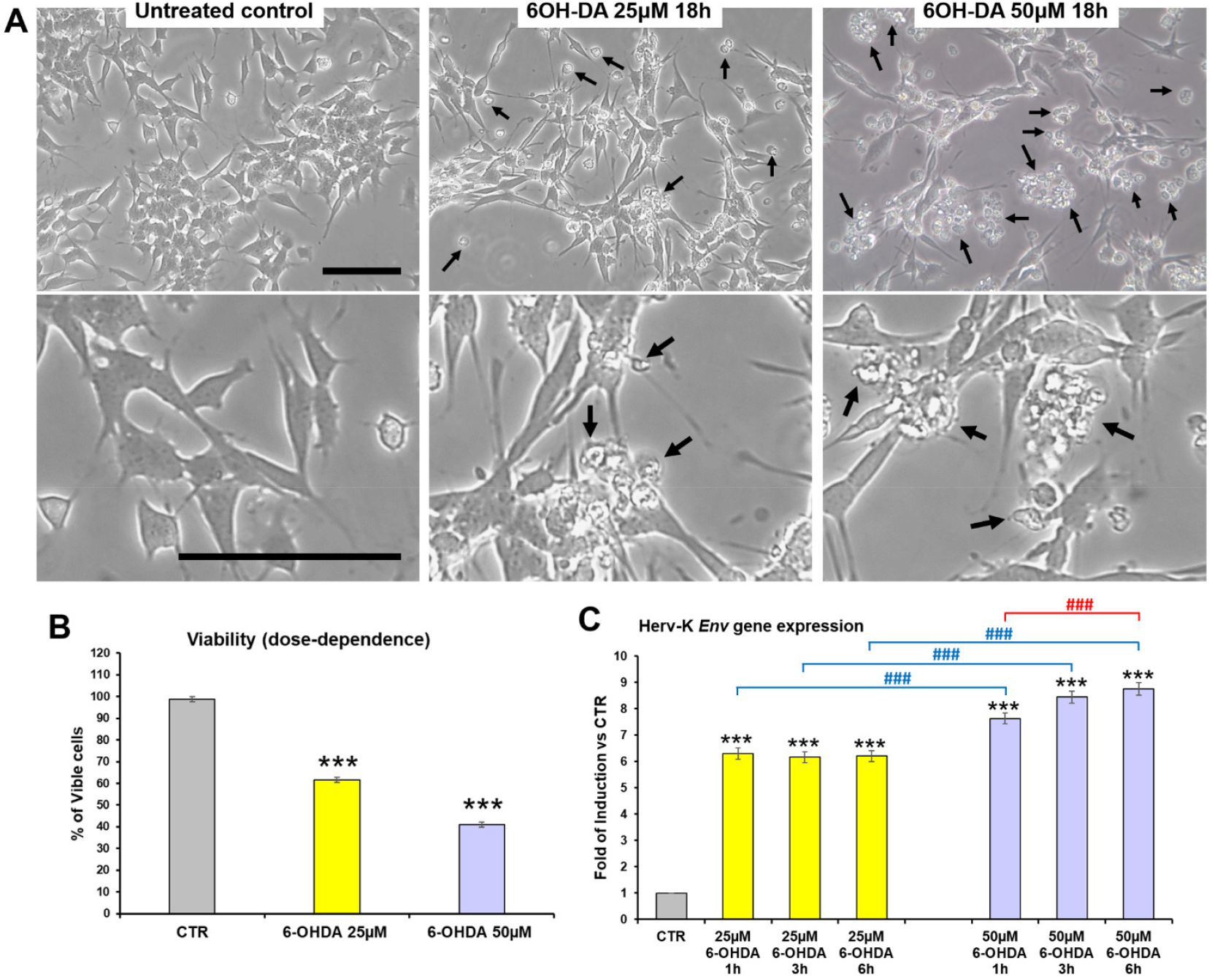
Dose-dependent neurotoxicity and early HERV-K activation induced by 6-OHDA in SHSY5Ywt cells. (**A**) Representative phase-contrast microscopy images of SHSY5Ywt cells exposed for 18 h to 25 µM or 50 µM 6-OHDA. Black arrows indicate morphological features of neurotoxic damage, including cell rounding, membrane blebbing, and cellular debris. Scale bars: 100 µm. (**B**) Quantitative analysis of cell viability following 18 h treatment with increasing concentrations of 6-OHDA, expressed as percentage of viable cells relative to control. (**C**) Time-course RT-qPCR analysis of HERV-K env gene expression following treatment with 25 µM or 50 µM 6-OHDA for 1 h, 3 h, and 6 h. Data are expressed as fold of induction vs control. Values represent mean ± SD from at least three independent experiments. ***p < 0.001 vs control; ###p < 0.001 vs 25 µM 6-OHDA at the same time point.

Consistent with these observations, quantitative analysis of cell viability demonstrated a significant, dose-dependent reduction in the percentage of viable cells, decreasing to approximately 60% and 40% of control levels following exposure to 25 µM and 50 µM 6-OHDA, respectively (**Fig. 4B**).

In parallel, RT-qPCR analysis revealed that 6-OHDA induced a rapid and robust transcriptional activation of the HERV-K Env sequence. Increased transcript levels were detected as early as 1 h after treatment and, for the higher dose, further increased at 3 h and 6 h. This response was dose-dependent, with 50 µM 6-OHDA eliciting a significantly stronger induction than 25 µM at all time points analyzed (**Fig. 4C**). Together, these results indicate that HERV-K de-repression represents an early molecular event associated with dopaminergic neurotoxicity in SHSY5Ywt cells.

### Early HERV-K activation is conserved in differentiated dopaminergic-like SHSY5Y cells and is associated with alterations in oxidative stress signaling

To determine whether HERV-K activation also occurs in a more neuronally differentiated context, RA-differentiated SHSY5Y cells (SHSY5Ydiff) were exposed to 6-OHDA. Phase-contrast microscopy revealed pronounced neurodegenerative changes already at early time points (3 h), which became more severe after 6 h of exposure to 75 µM 6-OHDA, including widespread neurite fragmentation and loss of cellular integrity (**Fig. 5A**).

**Figure 5.**
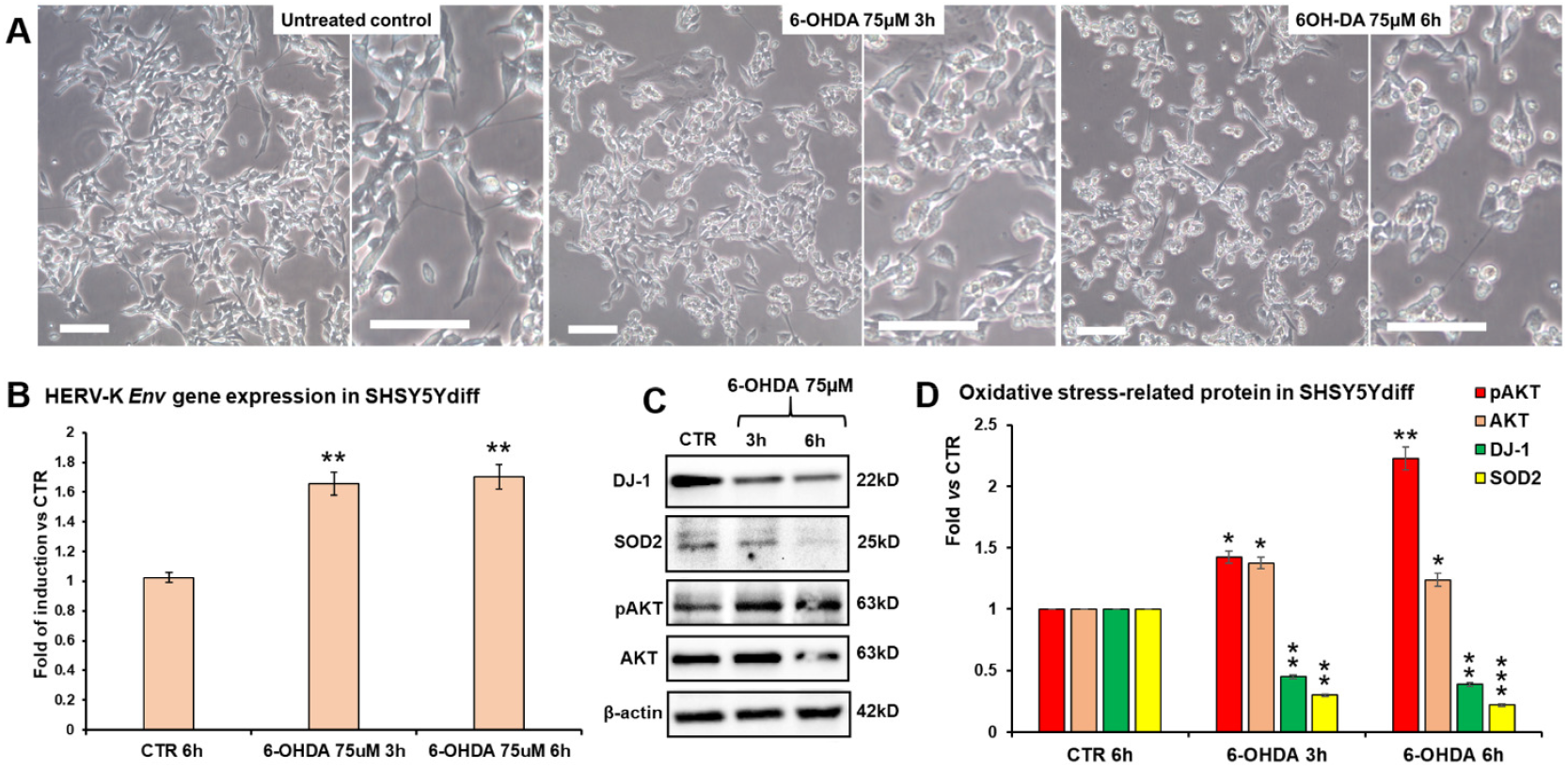
HERV-K activation and oxidative stress signaling alterations in differentiated SHSY5Y cells. (**A**) Representative phase-contrast microscopy images of retinoic acid–differentiated SHSY5Y cells (SHSY5Ydiff) under control conditions or following exposure to 75 µM 6-OHDA for 3 h or 6 h. Scale bars: 100 µm. (**B**) RT-qPCR analysis of HERV-K env gene expression in SHSY5Ydiff cells treated with 75 µM 6-OHDA for 3 h and 6 h, expressed as fold of induction vs control. (**C**) Representative Western blot analysis of oxidative stress–related and survival proteins (DJ-1, SOD2, pAKT, AKT) following 6-OHDA exposure. β-actin was used as a loading control. (**D**) Densitometric quantification of protein expression levels normalized to β-actin and expressed as fold change vs control. Data represent mean ± SD from at least three independent experiments.*p < 0.05, **p < 0.01 vs control.

RT-qPCR analysis demonstrated a significant induction of HERV-K Env expression in SHSY5Ydiff cells following neurotoxic stress, with transcript levels increased by approximately 1.6-1.8-fold at both 3 h and 6 h compared to untreated controls (**Fig. 5B**). These findings indicate that HERV-K de-repression is not restricted to proliferating cells but is preserved in differentiated dopaminergic-like neurons.

At the molecular level, 6-OHDA exposure was associated with marked alterations in oxidative stress-related and survival signaling pathways. WB analysis revealed a progressive reduction in the expression of the neuroprotective proteins DJ-1 and SOD2, together with dysregulation of AKT signaling, as reflected by changes in pAKT and total AKT levels (**Fig. 5C**). Densitometric analysis confirmed a significant decrease in antioxidant and survival markers concomitant with neurotoxic stress (**Fig. 5D**). These data suggest that early HERV-K activation occurs in parallel with oxidative stress responses and impairment of pro-survival signaling pathways.

### Modulation of the Wnt/β-catenin pathway attenuates 6-OHDA–induced HERV-K activation and oxidative stress responses

Given the established role of the Wnt/β-catenin pathway in neuronal survival and stress responses, we next investigated whether pharmacological modulation of this pathway influences HERV-K activation induced by neurotoxic stress. SHSY5Ywt cells were exposed to 6-OHDA in the presence or absence of atrial natriuretic peptide (ANP), which we previously demonstrated to be neuroprotective by modulating the Wnt/β-catenin signaling [35].

RT-qPCR analysis showed that pre-treatment with ANP significantly attenuated the 6-OHDA– induced upregulation of HERV-K *Env* transcripts at all time points examined. While 6-OHDA alone induced a strong, time-dependent increase in HERV-K expression, ANP co-treatment markedly reduced transcript levels, bringing them close to control values (**Fig. 6B**). Consistently, WB analysis demonstrated that ANP counteracted 6-OHDA–induced alterations in stress-related and survival proteins. ANP treatment restored the expression of antioxidant and neuroprotective markers, including SOD2, HSP70, DJ-1, and TH, and normalized AKT phosphorylation dynamics compared to cells treated with 6-OHDA alone (**Fig. 6C–F**). Quantitative analysis confirmed that ANP significantly mitigated oxidative stress signaling and preserved dopaminergic and survival markers in the context of neurotoxic insult.

**Figure 6.**
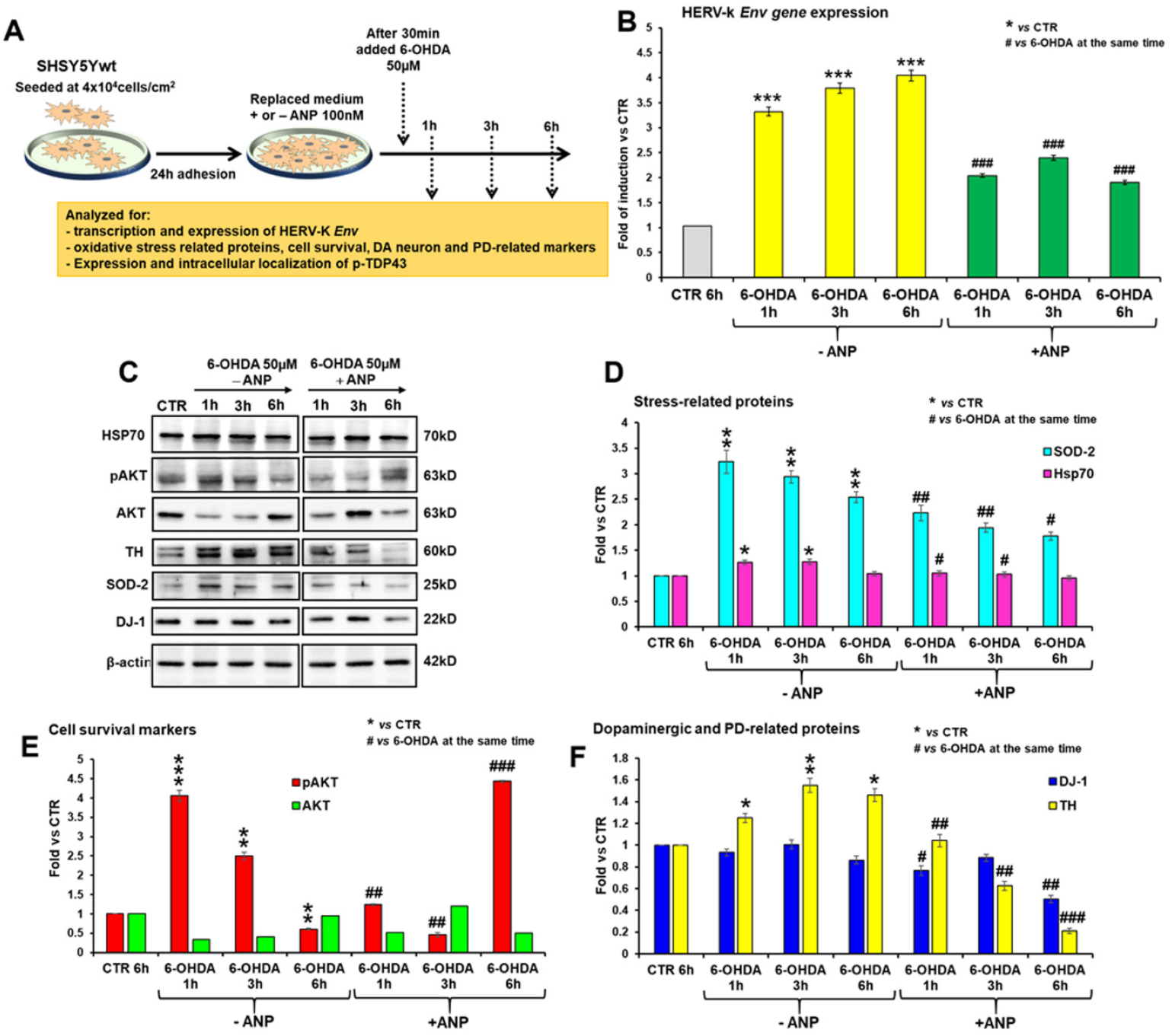
Modulation of Wnt/β-catenin signaling by ANP attenuates 6-OHDA–induced HERV-K activation and stress responses. (**A**) Experimental scheme illustrating SHSY5Ywt cell treatment protocol with 6-OHDA (50 µM) in the presence or absence of ANP (100 nM). **(B)** RT-qPCR analysis of HERV-K env gene expression at 1 h, 3 h, and 6 h following 6-OHDA exposure with or without ANP. Data are expressed as fold of induction *vs* control. (**C**) Representative Western blot analysis of stress-related, survival, and dopaminergic markers (HSP70, pAKT, AKT, TH, SOD2, DJ-1). (**D–F**) Densitometric quantification of stress-related proteins (SOD2, HSP70), survival markers (pAKT, AKT), and dopaminergic/PD-related proteins (DJ-1, TH), normalized to β-actin and expressed as fold change *vs* control. Data are presented as mean ± SD from at least three independent experiments. *p < 0.05, **p < 0.01 vs control; #p < 0.05, ##p < 0.01, ###p < 0.001 vs 6-OHDA alone at the same time point.

### Wnt/β-catenin modulation limits 6-OHDA–induced pTDP-43 accumulation and cytoplasmic mislocalization

Considering the reported neurotoxicity of the HERV-K Env protein [39], and the recent evidence indicating that activation of HERV-K expression is functionally linked to up-regulation and cytoplasmic accumulation of phosphorylated TAR DNA-binding protein 43 (pTDP-43) [26], we next investigated whether HERV-K over-transcription induced by neurotoxic stress was associated with increased Env protein levels as well as of pathological phosphorylation and mislocalization of TDP-43, and whether these processes might be modulated by ANP. WB and densitometric analyses revealed that 6-OHDA induced an early increase in both HERV-K Env protein levels and pTDP-43 in SHSY5Ywt cells, with pTDP-43 showing a non-monotonic temporal profile, characterized by a transient decrease at 3 h. Interestingly, ANP pre-treatment significantly reduced the accumulation of both proteins compared to cells treated with 6-OHDA alone. (**Fig. 7A–B**).

**Figure 7.**
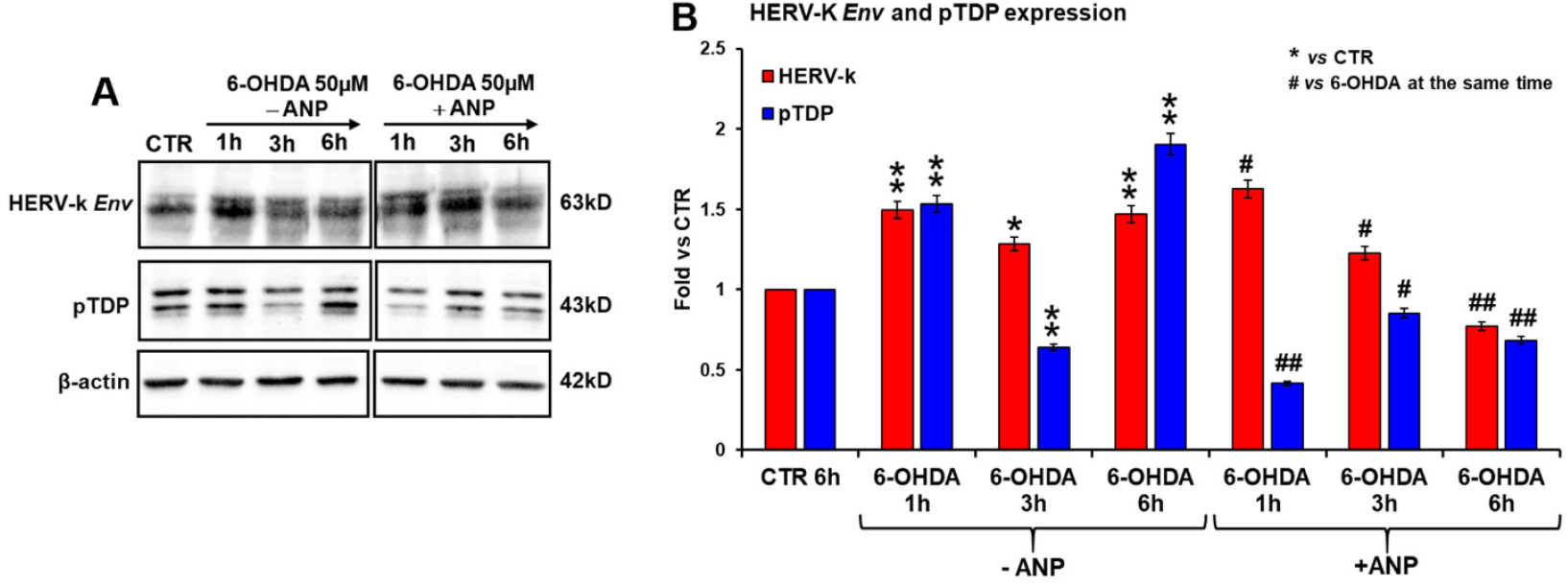
ANP reduces 6-OHDA–induced accumulation of HERV-K Env and phosphorylated TDP-43. (**A**) Representative Western blot analysis of HERV-K Env and phosphorylated TDP-43 (pTDP-43) in SHSY5Ywt cells treated with 50 µM 6-OHDA for 1 h, 3 h, or 6 h in the presence or absence of ANP (100 nM). β-actin was used as a loading control. (**B**) Densitometric quantification of HERV-K Env and pTDP-43 protein levels, normalized to β-actin and expressed as fold change vs control. Data represent mean ± SD from at least three independent experiments. *p < 0.05, **p < 0.01 vs control; #p < 0.05, ##p < 0.01 vs 6-OHDA at the same time point.

Immunofluorescence analysis further demonstrated that 6-OHDA exposure promoted a progressive cytoplasmic redistribution of pTDP-43, with a marked increase in the proportion of cells displaying cytoplasmic pTDP-43 aggregates over time. In contrast, ANP treatment significantly limited pTDP-43 mislocalization, preserving its predominantly nuclear localization (**Fig. 8A**). Quantitative analysis confirmed a substantial reduction in the percentage of cells positive for cytoplasmic pTDP-43 in the presence of ANP compared to 6-OHDA alone (**Fig. 8B**).

**Figure 8.**
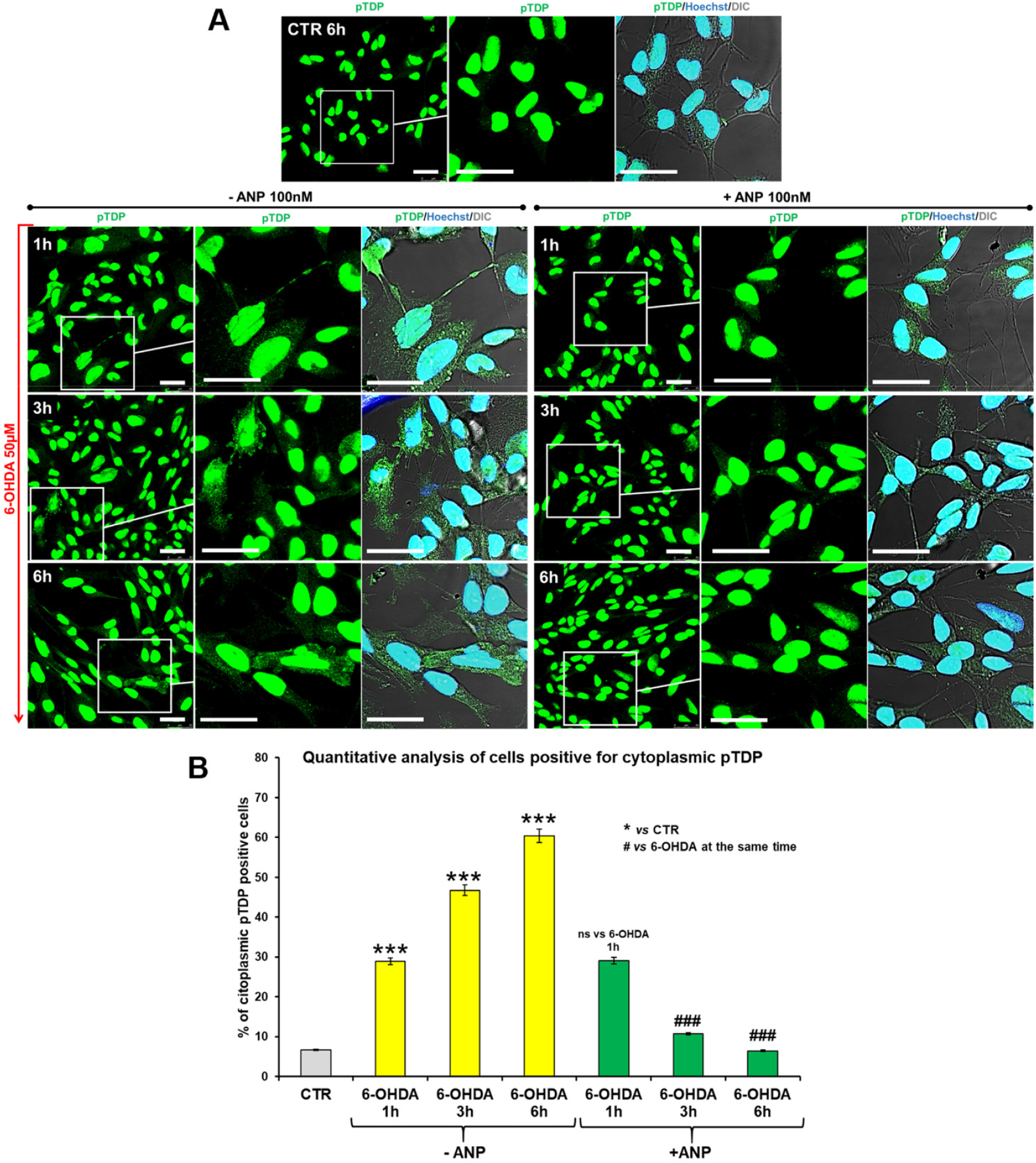
Wnt/β-catenin modulation prevents 6-OHDA–induced cytoplasmic mislocalization of pTDP-43. (**A**) Representative confocal microscopy images showing intracellular localization of phosphorylated TDP-43 (pTDP-43, green) in SHSY5Ywt cells treated with 50 µM 6-OHDA for 1 h, 3 h, or 6 h in the absence or presence of ANP (100 nM). Nuclei were counterstained with Hoechst (blue). Merged images include differential interference contrast (DIC). Insets highlight representative cells. Scale bars: 25 µm. (**B**) Quantitative analysis of the percentage of cells displaying cytoplasmic pTDP-43 localization. Data are expressed as mean ± SD from three independent experiments. ***p < 0.001 vs control; ###p < 0.001 vs 6-OHDA at the same time point; ns: not significant.

## DISCUSSION

In the present study, we investigated whether dopaminergic neurotoxic stress is associated with transcriptional activation of HERV-K sequences and whether modulation of pro-survival signaling pathways influences this response in PD-relevant cellular models. Using two complementary systems —undifferentiated SHSY5Y cells and DA neuron-like differentiated SHSY5Y cells — we show that exposure to the dopaminergic neurotoxin 6-OHDA induces a rapid and robust de-repression of HERV-K Env expression. This response occurs early after toxin exposure, scales with insult intensity, and is accompanied by alterations in oxidative stress defenses, survival signaling, and protein homeostasis. Notably, pharmacological modulation of the Wnt/β-catenin pathway by the natriuretic peptide ANP attenuates HERV-K activation and limits associated molecular alterations, including the accumulation and cytoplasmic mislocalization of phosphorylated TDP-43.

Endogenous retroelements are normally maintained in a transcriptionally repressed state through epigenetic mechanisms that preserve genomic stability and cellular homeostasis. However, pathological stress conditions — including oxidative stress, inflammation, and proteostatic imbalance — can relieve this repression [41, 42]. In NDs, ERV/HERV activation has been most extensively investigated in ALS and related proteinopathies, where it has been linked to neuroinflammation and neuronal toxicity [26, 43]. Although evidence in PD has been more limited, recent studies report altered HERV-K expression in PD-relevant cellular contexts, including astrocytes, suggesting that HERV-K dysregulation may contribute to disease-associated stress responses beyond motor neurons [44].

Within this framework, our data indicate that dopaminergic neurotoxic stress is sufficient to trigger HERV-K de-repression in neuronal models commonly used to study PD-related mechanisms. HERV-K Env induction was detectable within hours of 6-OHDA exposure, preceding overt cell death and persisting under conditions of reduced viability, supporting the idea that HERV-K activation represents an early stress-responsive event rather than a secondary consequence of terminal neurodegeneration. Similar early retroelement induction in response to oxidative or genotoxic stress has been reported in non-neuronal systems [42]. Oxidative stress and impaired redox homeostasis are central features of PD pathophysiology and are robustly reproduced in toxin-based cellular models [45]. In differentiated SHSY5Y cells, 6-OHDA exposure elicited coordinated alterations in oxidative stress–related proteins, including DJ-1 and SOD2, together with changes in AKT phosphorylation, concomitant with HERV-K activation. These findings suggest that redox imbalance and perturbed survival signaling may contribute to a permissive transcriptional environment for retroelement expression.

A key finding of this study is that ANP markedly counteracted neurotoxin-induced HERV-K activation and associated molecular changes. ANP has previously been shown to activate Wnt/β-catenin signaling and exert neuroprotective effects in dopaminergic cellular models [35, 36]. Given the pivotal role of Wnt/β-catenin signaling in neuronal survival and stress adaptation, and its dysregulation in PD [10], our data suggest that reinforcement of this pathway limits stress-driven HERV-K de-repression, potentially by restoring transcriptional or epigenetic constraints compromised under neurotoxic conditions.

The analysis of phosphorylated TDP-43 was motivated by recent evidence demonstrating a bidirectional relationship between ERV/HERV activation and TDP-43 proteinopathy [26, 43]. In addition, it has been reported that TDP-43 contributes to retrotransposon silencing and, particularly, directly regulates HERV-K transcription through binding to specific motifs within the HERV-K long terminal repeat (LTR), thereby contributing to the control of retroviral silencing under physiological conditions [43, 46]. Consistent with this framework, we observed that 6-OHDA increased both HERV-K Env and pTDP-43 levels and promoted cytoplasmic accumulation of pTDP-43, whereas ANP treatment reduced pTDP-43 accumulation and preserved its nuclear localization. Notably, Western blot analyses revealed a non-monotonic temporal profile of pTDP-43 levels following 6-OHDA exposure, characterized by a transient reduction at the intermediate time point, consistent with the dynamic and reversible regulation of TDP-43 phosphorylation under acute cellular stress [47]. Overall, these results support the notion that modulation of stress-responsive signaling pathways can dampen molecular interactions linking retroelement activation to proteostatic dysfunction.

From a translational perspective, HERV-K expression may represent an early molecular indicator of neurotoxic stress in PD-relevant models, reflecting upstream disturbances in redox balance and survival signaling. Moreover, strategies aimed at reinforcing pro-survival pathways such as Wnt/β-catenin signaling may provide dual benefits by supporting neuronal resilience while constraining retroelement activation and downstream proteinopathy-associated features. Although limited to 2D cellular systems, these findings are consistent with broader evidence implicating ERV/HERV dysregulation and Wnt/β-catenin signaling in ND mechanisms [10, 42].

On this ground, the use of molecules acting as modulators of the Wnt/β-catenin pathway—capable of exerting neuroprotective effects not only by targeting a signaling cascade whose dysregulation is implicated in neurodegenerative disorders, but also by directly or indirectly counteracting the reactivation of HERV-K retroviral sequences—may represent a promising strategy to halt the onset and progression of PD and other NDs [10, 31].

Some limitations should be acknowledged. Toxin-based cellular models capture key aspects of dopaminergic stress but do not reproduce the full cellular and circuit-level complexity of PD. Moreover, while we document coordinated changes in HERV-K expression, oxidative stress markers, survival signaling, and pTDP-43 expression and localization, direct causality among these processes remains to be established. Given the pleiotropic nature of ANP signaling, future studies using genetic approaches or alternative Wnt/β-catenin modulators will be required to confirm pathway specificity.

In conclusion, this study identifies HERV-K de-repression as an early stress-associated feature in PD-like cellular models and demonstrates that modulation of Wnt/β-catenin signaling by ANP limits HERV-K activation and associated molecular hallmarks, including oxidative stress responses and pTDP-43 pathology. These findings support the emerging concept that endogenous retroelements intersect with stress and survival pathways in neurodegeneration and highlight their potential relevance for mechanistic and therapeutic exploration in PD.

## ACKNOWLEDGEMENTS

This work was funded by Fondazione Intesa Sanpaolo “Fondo di Beneficenza” (project B/2023/0162 to A.S.), and supported by CNR (“InvAt-Invecchiamento Attivo e in Salute”, project DSB.AD006.371.001 to A.S.) and by European Union - Next Generation EU, PNRR project “Rome Technopole—Innovation Ecosystem” (to A.S. and S.R.).

## AUTHOR CONTRIBUTIONS

EKK, supervised the experiments, collected and analyzed data, and contributed to drafting the manuscript; SR and GDM performed experiments; SR supervised the RT-qPCR experiments; MC critically revised the manuscript; AS designed the study, supervised all experiments, analyzed the data, performed microscopic analyses, carried out the statistical analysis, and drafted the manuscript.

## DATA AVAILABILITY

All data generated in this study are included in this article, either as main figures or supplementary material. Primary data files are available from the corresponding author upon reasonable request.

