## Supplementary Materials for "Interplay Between Retroviral Element De-repression and Wnt/β-Catenin Pathway in Cellular Models of Parkinson’s Disease"

### SUPPLEMENTAL MATERIALS AND METHODS

#### Cell cultures and treatments

Exponentially growing SHSY5Y (SHSY5Ywt; Model A) cells were grown as a monolayer in Eagle's Minimum Essential Medium (α-MEM) plus HAM's F12 (1:1), supplemented with 10% heat-inactivated fetal bovine serum (FBS), L-glutamine (2 mM), penicillin (100 IU/ml), and streptomycin (100 µg/ml), and maintained at 37 °C, in a humidified atmosphere of 5% $CO_2$. For passaging, cells were detached from culture flasks with 0.05% trypsin and 0.002% EDTA solution. All media and supplements for cell cultures were acquired from Hyclone (Logan, UT, USA).

#### Western Blot (WB) Analysis

Cells were lysed in RIPA buffer containing 50 mM Tris-HCl (pH 8.0), 150 mM NaCl, 1% Nonidet NP-40, 0.5% sodium deoxycholate, 0.1% SDS, 1 mM PMSF, 2 mM $Na_3VO_4$, 20 mM NaF, and 1% protease inhibitor cocktail (Sigma-Aldrich, ST. Louis, MO, USA). Cell lysates were cleared by centrifugation, and protein concentrations were measured using the Bradford assay (Bio-Rad, Segrate, Italy). Equal amounts of protein (15 – 20 µg) were resolved by SDS–PAGE on 10% or 15% polyacrylamide gels, transferred onto nitrocellulose membranes (Hybond, Amersham GE Healthcare), blocked with 5% non-fat dry milk (Bio-Rad) in Tris-buffered saline containing 0.05% Tween-20 (TBS-T; 20 mM Tris, 150 mM NaCl, pH 7.6), and incubated with the primary antibodies reported and detailed in **Supplementary Table S1**. Primary antibody binding was detected using peroxidase-conjugated secondary antibodies (Bio-Rad). Membranes were visualized and images acquired using the ChemiDoc XRS+ imaging system (Bio-Rad). Band intensities were quantified using ImageJ (http://rsbweb.nih.gov/ij/). Densitometric values, normalized to β-actin, were expressed as uncalibrated optical density (OD) values or as fold change relative to untreated controls. Data represent at least three independent experiments and are reported as mean ± SD.

##### Quantitative Reverse Transcription Polymerase Chain Reaction (RT-qPCR)

For assessing the transcription levels of HERV-k Env and Gag genes by RT-qPCR, total RNA was extracted using TRIzol™ Reagent (Invitrogen, Carlsbad, CA, United States), and treated with RNase-free DNase Promega (Madison, WI, USA) according to the manufacturer's instructions. After extraction with a mixture of Phenol:Chloroform:Isoamyl Alcohol (25:24:1, Invitrogen) and precipitation with sodium acetate and ethanol, 1 µg of total RNA was reverse transcribed using random primers with ImProm-II Reverse Transcription system (Promega) according to the manufacturer's instructions. RT-qPCR was performed with iTaq Universal SYBR Green Supermix (Bio-Rad) using 350 nM of the specific primers, by the CFX Connect Real-Time PCR Detection System (Bio-Rad). Sequences of primers used are reported in **Supplementary Table S2**. The amplification conditions were: 1 min at 95 °C, followed by 40 cycles of 10 s at 95 °C and 40 s at 60 °C. Cq values were determined from the system software using 'single threshold' mode. The relative expression level of each gene was calculated from these Cqs using experimentally determined amplification efficiencies and then normalized to the reference gene GAPDH. Results were reported as fold of induction *vs* untreated control. In all experiments, each sample was analyzed in triplicate, and no-template controls and no-reverse transcription controls were included.

**Supplementary Table S1**. List of antibodies/reagents used for Western blot analyses.

| Antigen | Host | Cat. # | Working dilution | Supplier |
|---|---|---|---|---|
| DJ-1 (Parkinson disease protein 7) | Rabbit (polyclonal) | AB9212 | 1:5000 | (Burlington, MA, USA) |
| SOD2 (Superoxide Dismutase 2) | Mouse (monoclonal) | sc-133134 | 1:500 | Santa Cruz Biotechnology (Dallas, TX, USA) |
| AKT (protein kinase B) | Rabbit (polyclonal) | 9272 | 1:2000 | Cell signaling (Boston, MA, USA) |
| pAKT $^{Thr308}$ (active form) | Rabbit (polyclonal) | 9275 | 1:2000 | Cell signaling (Boston, MA, USA) |
| HSP70 (Heat Shock Protein 70) | Rabbit (polyclonal) | 4872 | 1:1000 | Cell signaling (Boston, MA, USA) |
| TH (Tyrosine hydroxylase) | Rabbit (polyclonal) | 2792 | 1:1000 | Cell signaling (Boston, MA, USA) |
| HERV-k *Env* (envelope protein) | Mouse (monoclonal) | HERM-1811-5 | 1:2000 | AMSBIO (Abingdon, UK; Cambridge, MA, USA) |
| pTDP-43 $^{Ser409/410}$ (TAR DNA-binding protein 43) | Rabbit (polyclonal) | 22309-1-AP | 1:2000 (WB) 1:200 (IF) | Proteintech (Rosemont, IL, USA) |
| β-actin | Mouse (monoclonal) | 4700 | 1:5000 | Sigma-Aldrich (St. Louis, MO, USA) |

**Supplementary Table S2**. Sequences of primers used quantitative reverse transcription polymerase chain reaction (RTqPCR)

| Gene Name | Forward Sequence | Reverse Sequence |
|---|---|---|
| HERV-k *Env* | CTGAGGCAATTGCAGGAGTT | GCTGTCTCTTCGGAGCTGTT |
| HERV-k *Gag* | AGCAGGTCAGGTGCCTGTAACATT | TGGTGCCGTAGGATTAAGTCTCCT |
| GAPDH | CCACATCGCTCAGACACCAT | ATGTAAACCATGTAGTTGAGG |

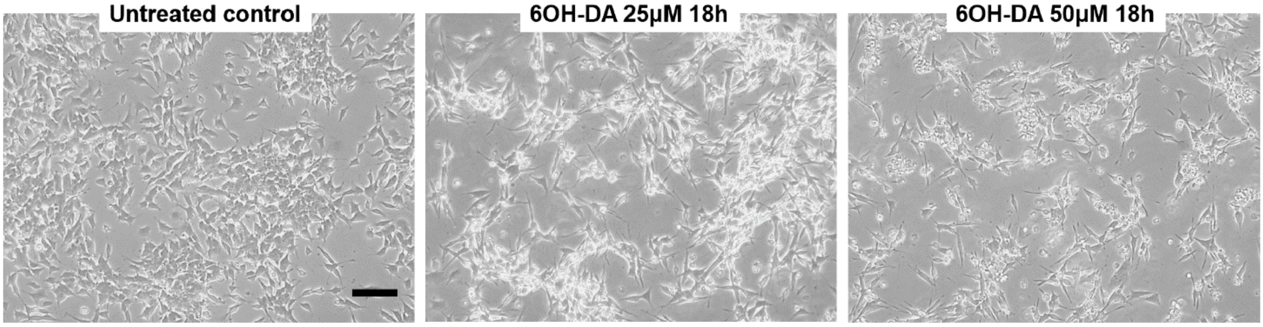


**Supplementary Figure S1. Dose-dependent neurotoxicity induced by 6-OHDA in SHSY5Ywt cells.** Representative low-magnification phase-contrast images showing the dose-dependent effects of 6-OHDA on SHSY5Y cell morphology and adhesion. Untreated cells display a dense and well-adherent monolayer, whereas exposure to increasing concentrations of 6-OHDA (25 µM and 50 µM for 18 h) results in progressive loss of cell–cell contacts, reduced adhesion, and marked morphological alterations. Images represent selected fields from the experimental conditions shown in Fig. 4A and are provided to illustrate the overall impact of neurotoxic stress on cell monolayer organization. Scale bar: 100 µm.
